# Dynamics of Spike-Specific Neutralizing Antibodies Across Five-Year Emerging SARS-CoV-2 Variants of Concern Reveal Conserved Epitopes that Protect Against Severe COVID-19

**DOI:** 10.1101/2024.09.22.614369

**Authors:** Latifa Zayou, Swayam Prakash, Hawa Vahed, Nisha Rajeswari Dhanushkodi, Afshana Quadiri, Ahmed Belmouden, Zohra Lemkhente, Aziz Chentoufi, Daniel Gil, Jeffrey B. Ulmer, Lbachir BenMohamed

## Abstract

Since early 2020, several SARS-CoV-2 variants of concern (VOCs) continue to emerge, evading waning antibody mediated immunity produced by the current Spike-alone based COVID-19 vaccines. This caused a prolonged and persistent COVID-19 pandemic that is going to enter its fifth year. Thus, the need remains for innovative next generation vaccines that would incorporate protective Spike-derived B-cell epitopes that resist immune evasion. Towards that goal, in this study we (*i*) Screened the sequences of Spike among many VOCs and identified conserved and non-conserved linear B-cell epitopes; (*ii*) Compared titers and neutralization antibodies specific to these conserved and non-conserved B-cell epitopes from serum of symptomatic and asymptomatic COVID-19 patients that were exposed to multiple VOCs across the 5^-^year COVID-19 pandemic, and (*iii*) Compared protective efficacy of conserved versus non-conserved B-cell epitopes against the most pathogenic Delta variant in a “humanized” ACE-2/HLA transgenic mouse model. We found robust conserved B-cell epitope-specific antibody titers and neutralization in sera from asymptomatic COVID-19 patients. In contrast, sera from symptomatic patients contained weaker antibody responses specific to conserved B-cell epitopes. A multi-epitope COVID-19 vaccine that incorporated the conserved B-cell epitopes, but not the non-conserved B-cell epitopes, significantly protected the ACE2/HLA transgenic mice against infection and COVID-19 like symptoms caused by the Delta variant. These findings underscore the importance of conserved B-cell epitopes in generating robust protective immunity against severe COVID-19 symptoms caused by various VOCs, providing valuable insights for the development of broad-spectrum next generation Coronavirus vaccines capable of conferring cross-variant protective immunity.

**IMPORTANCE:** A persistent COVID-19 pandemic continues to evolve because of a continued emergence of SARS-CoV-2 variants of concern (VOCs) that escape the antibodies induced by the current Spike-alone COVID-19 vaccines. Identifying and characterizing the protective and non-protective Spike-derived B-cell epitopes that resist immune-evasion is a paramount for the development of broad-spectrum next generation Coronavirus vaccines. The present study identified Spike-derived conserved B cell epitopes that (*i*) are targeted by consistent and strong antibody responses in asymptomatic COVID-19 patients across the 5-year pandemic regardless of VOCs; and (*ii*) provided strong protection in ‘humanized” ACE2/HLA transgenic mice against infection and COVID-19 like symptoms caused by the most pathogenic Delta variant. The findings have the potential to inform the design of next generation Coronavirus vaccines capable of conferring cross-variant protective immunity.

**TWEET:** Protective SARS-CoV-2 Conserved Linear B Cell Epitopes Identified from Spike Protein.

## INTRODUCTION

Since early 2020, the world has encountered successive waves of COVID-19, fueled by the emergence of over 20 variants of concern (VOCs) with maintained transmissibility and virulence (4). As of September 2024, the number of confirmed SARS-CoV-2 cases has reached over 776 million, and COVID-19 has caused over 7 million deaths (1–3). The world will enter its sixth year of a persistent COVID-19 pandemic, fueled by the continuous emergence of heavily Spike-mutated and highly contagious SARS-CoV-2 variants and sub-variants that continue to: (*i*) escaped immunity induced by the current Spike-alone-based vaccines; (*ii*) disrupt the efficacy of the COVID-19 booster paradigm (5–10); and (*iii*) outpace the development of variant-adapted bivalent Spike-alone vaccines (1–3, 11, 12). This bleak outlook of a prolonged COVID-19 pandemic emphasizes the urgent need for developing a next-generation broad-spectrum pan-Coronavirus vaccine capable of conferring strong cross-variant and cross-strain protective immunity that would prevent immune evasion and breakthrough infections (12).

The Spike protein is heavily mutated in these variants with an accumulated 346 mutations since the ancestral Wuhan strain, including 60 and 52 new mutations, in BA.2.86 and JN.1 Omicron subvariants, respectively. As such the Omicron variant exhibits reduced susceptibility to vaccine-induced neutralizing antibodies, requiring a boost to generate protective immunity (13). The dynamics of SARS-CoV-2 virus neutralization are integral to understanding and enhancing the efficacy of COVID-19 vaccines. Understanding the dynamics of Spike neutralizing antibodies would inform vaccine booster strategies and help to predict the impact of new variants on vaccine-induced immunity. The changing dynamics of SARS-CoV-2 virus neutralization plays a crucial role in determining the efficacy of COVID-19 vaccines and it depends on (***i***) the level of Neutralizing Antibody production, (***ii***) the role of emerging SARS-Cov-2 variants of concerns (VOCs), (***iii***) the role of immune escape mechanisms by emerging SARS-CoV-2 variants, and (***iv***) booster doses of COVID-19 vaccines. We previously, mapped and characterized the antigenicity and immunogenicity of genome-wide linear B cell epitopes that are highly conserved (1). We demonstrated that conserved B cell epitopes provided cross-protection against all the emerging SARS-CoV-2 variants of concern (VOCs), i.e., Alpha (B.1.1.7), Beta (B.1.351), Gamma (P.1), Epsilon (B.1.427/B.1.429), Delta (B.1.617.2), and Omicron (B.1.1.529) in a north American COVID-19 population cohort. In the present study, we hypothesize that multi-epitope vaccine candidates that incorporate highly conserved, antigenic, and immunogenic B cell epitopes will provide broader protection against COVID-19 caused by multiple SARS-CoV-2 VOCs.

We identified six B cell epitopes that are highly conserved within all the 20 VOCs of SARS-CoV-2, SARS-CoV, MERS-CoV, common cold Coronaviruses (HKU, OC1, 229E, NL63), and animal CoVs strains (i.e., Bats, Civet Cats, Pangolin and Camels). We established that those epitopes were selectively recognized by antibodies from “naturally protected” asymptomatic COVID-19 patients. In contrast symptomatic patients exhibited weaker antibody titers specific to conserved B-cell epitopes. Accordingly, antibodies from asymptomatic COVID-19 patients, but not from symptomatic patients, exhibited higher neutralization efficacy. Immunization of ACE2/HLA transgenic mice with a mixture of conserved B-cell epitopes significantly protected against infection and COVID-19 like symptoms caused by Delta variant. This study underscores the importance of conserved B-cell epitopes in generating robust protective immunity against severe disease caused by various SARS-CoV-2 variants, providing valuable insights for the development of broad-spectrum vaccines against COVID-19.

## MATERIALS & METHODS

### Human study population cohort and HLA genotyping

We enrolled 210 subjects from a pool of over 682 subjects. Written informed consent was obtained from participants before inclusion. The subjects were categorized as mild to severe COVID-19 groups and have undergone treatment at the University of California Irvine Medical Center between July 2020 to July 2022 (Institutional Review Board protocol # 2020-5779). SARS-CoV-2 positivity was defined by a positive RT-PCR on nasopharyngeal swab samples. All the subjects were genotyped by PCR for class I HLA-A*02:01 and class II HLA-DRB1*01:01 among the 682 patients (and after excluding a few for which the given amount of blood was insufficient - i.e., less than 6ml), ending with 210 that were genotyped for HLA-A*02:01^+^ or/and HLA-DRB1*01:01^+^ (15, 16). Based on the severity of symptoms and ICU admission/intubation status, the subjects were divided into two broad severity categories namely: Symptomatic (patients who died from COVID-19 complications, infected COVID-19 patients with severe disease that were admitted to the intensive care unit (ICU) and required ventilation support, infected COVID-19 patients with severe disease that required enrollment in ICU but without ventilation support, infected COVID-19 patients with moderate symptoms that involved a regular hospital admission or infected COVID-19 patients with mild symptoms) and Asymptomatic (infected individuals with no symptoms).

### Sequence comparison among variants of SARS-CoV-2 and animal CoV strains

We retrieved nearly 8.5 million human SARS-CoV-2 genome sequences from the GISAID database representing countries from North America, South America, Central America, Europe, Asia, Oceania, Australia, and Africa. All the sequences included in this study were retrieved either from the NCBI GenBank (www.ncbi.nlm.nih.gov/nuccore) or GISAID (www.gisaid.org). Multiple sequence alignment was performed keeping SARS-CoV-2-Wuhan-Hu-1 (MN908947.3) protein sequence as a reference against all the SARS-CoV-2 VOCs, common cold, and animal CoV strains. The sequences were aligned using the high throughput alignment tool DIAMOND (17).This comprised of all the VOCs and VBMs of SARS-CoV-2 (B.1.177, B.1.160, B.1.1.7, B.1.351, P.1, B.1.427/B.1.429, B.1.258, B.1.221, B.1.367, B.1.1.277, B.1.1.302, B.1.525, B.1.526, S:677H.Robin1, S:677P.Pelican, B.1.617.1, B.1.617.2, B,1,1,529) and common cold SARS-CoV strains (SARS-CoV-2-Wuhan-Hu-1 (MN908947.3), SARS-CoV-Urbani (AY278741.1), HKU1-Genotype B (AY884001), CoV-OC43 (KF923903), CoV-NL63 (NC_005831), CoV-229E (KY983587)) and MERS (NC_019843)). We have included whole-genome sequences from the bat ((RATG13 (MN996532.2), ZXC21 (MG772934.1), YN01 (EPI_ISL_412976), YN02(EPI_ISL_412977), WIV16 (KT444582.1), WIV1 (KF367457.1), YNLF_31C (KP886808.1), Rs672 (FJ588686.1)), pangolin (GX-P2V (MT072864.1), GX-P5E (MT040336.1), GX-P5L (MT040335.1), GX-P1E (MT040334.1), GX-P4L (MT040333.1), GX-P3B (MT072865.1), MP789 (MT121216.1), Guangdong-P2S (EPI_ISL_410544)), camel (KT368891.1, MN514967.1, KF917527.1, NC_028752.1), and civet (Civet007, A022, B039)) in evaluating the evolutionary relationship among the SARS-CoV-2 variants and common cold CoV strains,

### Data and Code Availability

The human-specific SARS-CoV-2 complete genome sequences were retrieved from the GISAID database, whereas the SARS-CoV-2 sequences for pangolin (*Manis javanica*), and bat (*Rhinolophus affinis*, *Rhinolophus malayanus*) were retrieved from NCBI. Genome sequences of previous strains of SARS-CoV for humans, bats, civet cats, and camels were retrieved from the NCBI GenBank.

### SARS-CoV-2 B Cell Epitope Prediction

Linear B cell epitope predictions were carried out on the spike glycoprotein (S), the primary target of B cell immune responses for SARS-CoV. We used the BepiPred 2.0 algorithm embedded in the B cell prediction analysis tool hosted on the IEDB platform. For each protein, the epitope probability score for each amino acid and the probability of exposure was retrieved. Potential B cell epitopes were predicted using a cutoff of 0.55 (corresponding to a specificity greater than 0.81 and sensitivity below 0.3) and considering sequences having more than 5 amino acid residues. This screening process resulted in 8 B-cell peptides. These epitopes represent all the major non-synonymous mutations reported among the SARS-CoV-2 variants. One B-cell epitope (S_439-482_) was observed to possess the maximum number of variant-specific mutations. Structure-based antibody prediction was performed using Discotope 2.0, and a positivity cutoff greater than −2.5 was applied (corresponding to specificity greater than or equal to 0.80 and sensitivity below 0.39), using the SARS-CoV-2 spike glycoprotein structure (PDB ID: 6M1D).

### Protein-peptide molecular docking

Computational peptide docking of B cell peptides into the ACE2 complex (binding protein) was performed using the GalaxyPepDock under GalaxyWEB. To retrieve the ACE2 structure, we used the x-ray crystallographic structure ACE2-B0AT1 complex 6M1D available on the Protein Data Bank. The 6M1D with a structural weight of 334.09 kDa possesses two unique protein chains, 2,706 residues, and 21,776 atoms. In this study, flexible target docking based on an energy-optimization algorithm was carried out on the ligand-binding domain containing ACE2 within the 4GBX structure. Similarity scores were calculated for protein-peptide interaction pairs for each residue. The prediction accuracy is estimated from a linear model as the relationship between the fraction of correctly predicted binding site residues and the template-target similarity measured by the protein structure similarity score and interaction similarity (S_Inter_) score obtained by linear regression. S_Inter_ shows the similarity of amino acids of the B cell peptides aligned to the contacting residues in the amino acids of the ACE2 template structure. Higher S_Inter_ score represents a more significant binding affinity among the ACE2 molecule and B cell peptides. Subsequently, molecular docking models were built based on distance restraints for protein peptide pairs using GalaxyPepDock. Based on the optimized energy scores, docking models were ranked.

### Peptide synthesis

Potential B cell epitopes identified from human-SARS-CoV-2 spike protein, were synthesized using solid-phase peptide synthesis and standard 9-fluorenylmethoxycarbonyl technology (21st Century Biochemicals, Marlborough, MA). The purity of peptides was over 90%, as determined by reversed-phase HPLC (Vydac C18) and mass spectroscopy (Voyager MALDI-TOF System). Stock solutions were made at 1 mg/ml in 10% DMSO in PBS.

### TaqMan quantitative polymerase reaction assay for the screening of SARS-CoV-2 Variants in COVID-19 patients

We utilized a laboratory-developed modification of the CDC SARS-CoV-2 RT-PCR assay, which received Emergency Use Authorization by the FDA on April 17th, 2020. (https://www.fda.gov/media/137424/download [accessed 24 March 2021]).

### Mutation screening assays

SARS-CoV-2-positive samples were screened by four multiplex RT-PCR assays. Through the qRT-PCR, we screened for 11 variants of SARS-CoV-2 in our patient cohort. The variants which were screened include B.1.1.7 (Alpha), B.1.351 (Beta), P.1 (Gamma), and B.1.427/B.1.429 (Epsilon), B.1.525 (Eta), R.1, P.2 (Zeta), B.1.526 (Iota), B.1.2/501Y or B.1.1.165, B.1.1.529 (BA.1) (Omicron), B.1.1.529 (BA.2) (Omicron), and B.1.617.2 (Delta). The sequences for the detection of Δ69–70 were adapted from a multiplex real-time RT-PCR assay for the detection of SARS-CoV-2 (18). The probe overlaps with the sequences that contain amino acids 69 to 70; therefore, a negative result for this assay predicts the presence of deletion S-Δ69–70 in the sample. Using a similar strategy, a primer/probe set that targets the deletion S-Δ242–244 was designed and was run in the same reaction with S-Δ69-70. In addition, three separate assays were designed to detect spike mutations S-501Y, S-484K, and S-452R and wild-type positions S-501N, S-484E, and S-452L.

Briefly, 5 µl of the total nucleic acid eluate was added to a 20 µl total-volume reaction mixture (1x TaqPath 1-Step RT-qPCR Master Mix, CG [Thermo Fisher Scientific, Waltham, MA], with 0.9 *m*M each primer and 0.2 *m*M each probe). The RT-PCR was carried out using the ABI StepOnePlus thermocycler (Life Technologies, Grand Island, NY). The S-N501Y, S-E484K, and S-L452R assays were carried out under the following conditions: 25°C for 2 min, then 50°C for 15 min, followed by 10 min at 95°C and 45 cycles of 95°C for 15 s and 65°C for 1 min. The Δ 69–70 / Δ242– 244 assays were run under the following conditions: 25°C for 2 min, then 50°C for 15 min, followed by 10 min at 95°C and 45 cycles of 95°C for 15 s and 60°C for 1 min. Samples displaying typical amplification curves above the threshold were considered positive. Samples that yielded a negative result or results in the S-Δ69–70/Δ242–244 assays or were positive for S-501Y P2, S-484K P2, and S-452R P2 were considered screen positive and assigned to a VOC.

### Enzyme-linked immunosorbent assay (ELISA)

Serum antibodies specific for epitope peptides and SARS-CoV-2 proteins were detected by ELISA. 96-well plates (Dynex Technologies, Chantilly, VA) were coated with 0.5 μg peptides or 100 ng S protein per well at 4°C overnight, respectively, and then washed three times with PBS and blocked with 3% BSA (in 1 X PBS) for 2hours. at RT. After blocking, the plates were incubated with 1:200 dilutions of the sera (100 μl/well) overnight at 4°C. The bound serum antibodies were detected with HRP-conjugated goat anti-mouse IgG and chromogenic substrate TMB (ThermoFisher, Waltham, MA). The cut-off for seropositivity was set as the mean value plus three standard deviations (3SD) in HBc-S control sera. The binding of the epitopes to the sera of SARS-CoV-2 infected samples was detected by ELISA using the same procedure, 96-well plates were coated with 0.5 μg peptides and sera were diluted at 1:50. All ELISA studies were performed at least twice.

### Neutralizing antibody assays for SARS-CoV-2

Serially diluted heat-inactivated plasma (1:3) and 300 PFU of SARS-CoV-2 variants were combined in Dulbecco’s Modified Eagle’s Medium (DMEM) and incubated at 37°C 5% CO_2_ for 30 minutes. After neutralization, the antibody-virus inoculum was transferred onto Vero E6 cells (ATCC C1008) and incubated at 34°C 5% CO_2_ for 1 hour. The cells were then fixed with 10% neutral buffered formalin and incubated at −20°C for 10 minutes followed by 20 minutes at room temperature. Plates were developed with True Blue HRP substrate and imaged on an ELISpot reader. The half maximum inhibitory concentration (IC50) was calculated using normalized counted foci.

### Triple transgenic mice immunization with multi-epitope Pan-Coronavirus vaccine and infection

The animal studies were performed at the University of California Irvine and adhered to the Guide for the Care and Use of Laboratory Animals published by the US National Institute of Health. All animal experiments were performed under the approved IACUC protocol # AUP-22-086. Female HLA-DR*0101/HLA-A*0201/hACE2 triple transgenic mice (8-9 weeks old) were used in this study. The HLA-DR*0101/HLA-A*0201/hACE2 triple transgenic mouse colony was established here at the UCI by cross-breeding K18-hACE2 mice (17) with double transgenic HLA-DR*0101/HLA-A*0201 mice (14).

The HLA-DR*0101/HLA-A*0201/hACE2 triple transgenic mice were immunized intranasally on day 0 with the multi-epitope coronavirus vaccine at 2 × 10^10^ viral particles [VP] per mouse, *n* = 35. The vaccine comprised of highly conserved and immunogenic 9 B cell epitopes, 16 CD8^+^ T cell epitopes, and 6 CD4^+^ T cell epitope. Fifteen mice were divided into 3 groups of 5 mice each including the multi-epitope vaccine group, control vaccine group and as a negative control, and the third group of 5 mice received sterile PBS (mock vaccinated group). The triple transgenic mice were intranasally infected with 1 x 10^4^ pfu of SARS-CoV-2 (Delta) delivered in 20 µL sterile PBS on day 28 following immunization (**Fig. 5A**). Mice were monitored daily for death and weight loss to day 14 post-infection (p.i.) on which they were euthanized for virological and immunological studies.

### Virus titration in oropharyngeal swabs

Throat swabs were analyzed for SARS-CoV-2 specific RNA by qRT-PCR. As recommended by the CDC, we used *ORF1ab-*specific primers (forward-5′-*CCCTGTGGGTTTTACACTTAA*-3′ and reverse-5′-*ACGATTGTGCATCAGCTGA*-3′) and probe (6FAM-*CCGTCTGCGGTATGTGGAAAGGTTATGG*-BHQ) to detect the viral RNA level in lungs. Briefly, 5 mL of the total nucleic acid eluate was added to a 20-mL total volume reaction mixture [1× TaqPath 1-Step RT-qPCR Master Mix (Thermo Fisher Scientific, Waltham, MA)], with 0.9 mM each primer and 0.2 mM each probe. The qRT-PCR was carried out using the ABI StepOnePlus thermocycler (Life Technologies, Grand Island, NY). When the Ct-value was relatively high (35 ≤ Ct < 40), the specimen was retested twice and considered positive if the Ct-value of any retest was less than 35.

### Statistical analyses

Data for each differentially expressed markers among blockade-treated and mock-treated groups of HLA-Tg mice were compared by ANOVA and Student t test using GraphPad Prism version 6 (GraphPad Software, La Jolla, CA). Statistical differences observed in the measured Ab responses between healthy donors and COVID-19 patients were calculated using ANOVA and multiple t test comparison procedures in GraphPad Prism. Data are expressed as the mean ± SD. Results were considered statistically significant at P < 0.05.

## RESULTS

### 1. Highly conserved B-cell epitopes identified in different SARS-CoV-2 variants of concerns

A total of 210 COVID-19 patients participated in the study, categorized into asymptomatic and symptomatic groups based on clinical parameters. Blood and nasopharyngeal swabs were collected from all subjects for further analysis. Utilizing a qRT-PCR assay, viral haplotypes unique to different SARS-CoV-2 VOCs were identified. Notably, six novel nonsynonymous mutations (Δ69-70, Δ242-244, N501Y, E484K, L452R, and T478K) were employed to differentiate variants including Omicron (B.1.1.529 (BA.1)), Omicron (B.1.1.529 (BA.2)), Alpha (B.1.1.7), Beta (B.1.351), Gamma (P.1), Delta (B.1.617.2), and Epsilon (B.1.427/B.1.429). Serum samples from COVID-19 patients infected with highly pathogenic SARS-CoV-2 VOCs were analyzed for anti-SARS-CoV-2 peptide-specific IgG levels. Graphical representation of optical density revealed the magnitude of IgG response, with dotted lines shown in the graph serve to highlight peptides that consistently exhibit high immunogenicity and stable responses across different SARS-CoV-2 variants of concern, when compared to other conserved peptides (**Fig. S1**). These dotted lines are used to visually emphasize the peptides that elicit stronger and more consistent IgG responses over time, despite the emergence of diverse viral variants compared to their counterparts, providing a clear indication of their reliability and effectiveness in eliciting immune responses. Among the 17 tested peptides (**Supplemental Table 1**), six conserved epitopes (S_287-317_, S_369-393_, S_471-501_, S_565-598_, S_614-640_, S_1133-1160_) were identified as highly immunogenic peptides. These six conserved epitopes exhibited robust immunogenicity and demonstrated conservation across multiple SARS-CoV-2 variants (**Fig. S1**). Sequence homology analysis was conducted to determine the conservancy of immunodominant B-cell epitopes among SARS-CoV-2 variants of concern (**Table. 1**). In parallel, IgG response to conserved (S_565-598_ and S_287-317_), and non-conserved (S_13-37_ and S_601-628_) epitopes was evaluated in COVID-19 patients infected with various SARS-CoV-2 variants demonstrated in the pie charts (**Fig. 3**). The analysis revealed that over time, the immunogenicity of conserved epitopes remained stable and higher compared to non-conserved epitopes. Conversely, the immunogenicity of non-conserved epitopes exhibited a declining trend over time (**Fig. 3**).

**Table 1.** SARS-CoV-2 variant screening in COVID-19 patients: The table represents the distribution of SARS-CoV-2 variants detected in 198 COVID-19 positive patients using TaqMan quantitative polymerase reaction (qRT-PCR) assays. Variants were identified based on 6 specific nonsynonymous mutations associated with each variant, including Δ69–70, Δ242–244, S-501Y, S-484K, and S-452R, among others. Samples showing positive results for variant-specific mutations highlighted in black were considered screen positive and assigned to the respective variant group.

|  | Nonsynonymous Mutation |  |  |  |  |  |  |
| --- | --- | --- | --- | --- | --- | --- | --- |
| Variants of Concern | Δ69-70 | 242-244 | N501Y | E484K | L452R | T478K | Total Sample Size (%) |
| B.1.1.7 (Alpha) | Positive | Negative | Positive | Negative | Negative | Negative | 22 (11.1) |
| B.1.351( Beta) | Negative | Positive | Positive | Positive | Negative | Negative | 17 (8.6) |
| P.1. (Gamma) | Negative | Negative | Positive | Positive | Negative | Negative | 19 (9.6) |
| B.1.617.2 (Delta) | Negative | Negative | Negative | Negative | Positive | Positive | 44 (22.2) |
| B.1.427/B.1.429 (Epsilon) | Negative | Negative | Negative | Negative | Positive | Negative | 31 (15.7) |
| B.1.1.529 (BA.1) Omicron | Positive | Negative | Positive | Positive | Negative | Positive | 35 (17.7) |
| B.1.1.529 (BA.2) Omicron | Negative | Negative | Positive | Positive | Negative | Positive | 30 (15.1) |

**Figure 1.**
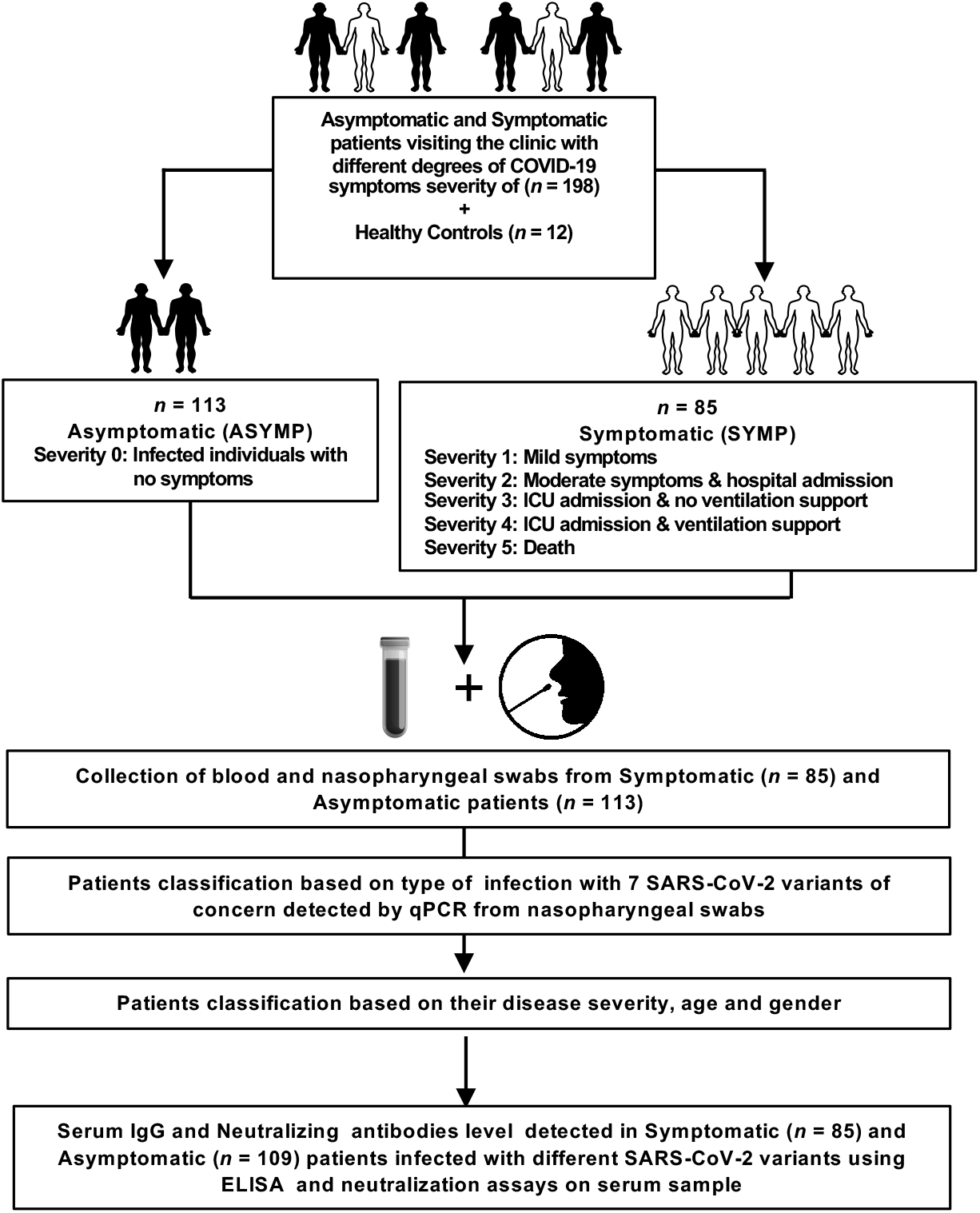
Assessment of SARS-CoV-2 infection severity and immune response: Experimental plan outlines the assessment of SARS-CoV-2 infection severity and immune response in a cohort of 198 patients, comprising both asymptomatic and symptomatic individuals presenting to the clinic with varying degrees of COVID-19 symptoms. Additionally, 12 healthy controls are included for comparative analysis. The patient cohort is categorized into asymptomatic (ASYMP) and symptomatic (SYMP) groups, with severity levels ranging from 0 to 5 based on symptom severity and clinical outcomes. Asymptomatic individuals exhibit no symptoms, while symptomatic patients are further classified based on the severity of their symptoms, hospital admission, ICU admission, ventilation support, and mortality. Blood and nasopharyngeal swabs are collected from both symptomatic (n = 85) and asymptomatic (n = 113) patients for subsequent analysis. Patients are classified based on the detection of 7 SARS-CoV-2 variants of concern using qPCR from nasopharyngeal swabs, as well as their disease severity, age, and gender. Serum IgG and neutralizing antibody levels are measured in symptomatic (n = 85), and asymptomatic (n = 109) patients infected with different SARS-CoV-2 variants using enzyme-linked immunosorbent assay (ELISA) and neutralization assays on serum samples.

**Figure 2.**
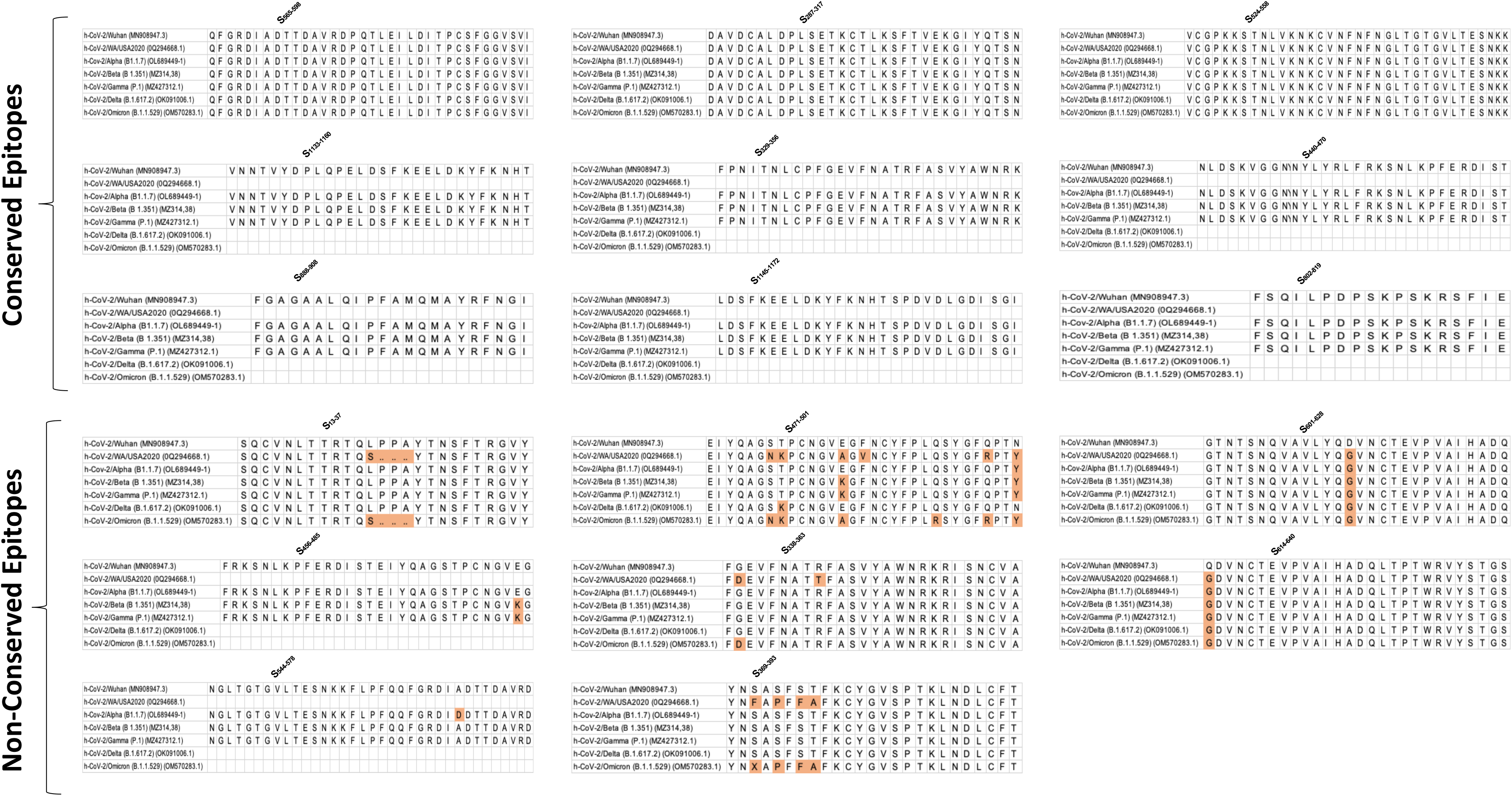
Sequence homology analysis of immunodominant B cell epitopes among SARS-CoV-2 variants of concern: The figure represents the results of sequence homology analysis to assess the degree of conservancy of immunodominant B cell epitopes among SARS-CoV-2 variants of concern. Seventeen peptides, identified as potential B cell epitopes, were subjected to analysis, and categorized into conserved epitopes and non-conserved epitopes based on sequence similarity across different variants. Conserved epitopes exhibit a high degree of sequence conservation among variants, suggesting potential cross-reactivity and broad immune recognition. Non-conserved epitopes, on the other hand, show variability in sequence composition across variants, indicating potential immune evasion and reduced recognition by antibodies.

**Figure 3.**
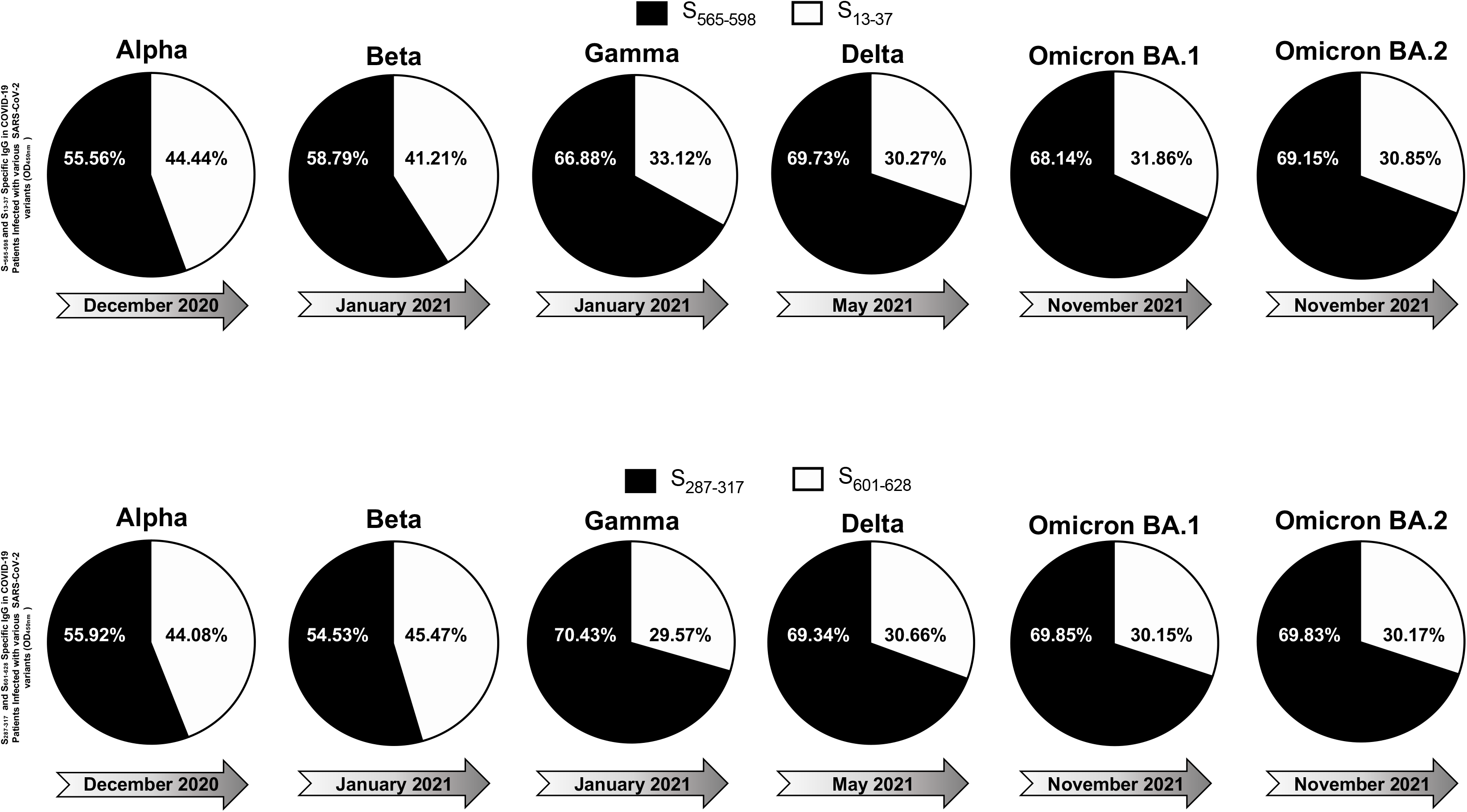
IgG Response to conserved and non-conserved epitopes in COVID-19 patients infected with different SARS-CoV-2 variants: The figure displays six pie charts representing the IgG response to S_565-598_ and S_13-37_ epitopes (upper panel) and S_287-317_ and S_601-628_ (bottom panel) in COVID-19 patients infected with various SARS-CoV-2 variants of concern. Each pie chart corresponds to a specific variant, including Alpha, Beta, Gamma, Delta, Omicron BA.1, and Omicron BA.2, which appeared at different time points during the pandemic. In each pie chart, black segments represent the IgG response against conserved peptide sequences (S_565-598_ and S_287-317_), while white segments represent the IgG response against non-conserved peptide sequences (S_13-37_ and S_601-628_). The intensity of the black and white segments indicates the relative magnitude of the IgG response to conserved and non-conserved epitopes, respectively.

### 2. Severity, age, and gender-dependent antibody responses to conserved B-cell epitopes in COVID-19 patients

The immunogenicity of ‘universal’ B-cell epitopes in COVID-19 patients was examined across different severity groups, age categories, and genders, with a focus on humoral immune responses. ELISA assays were conducted to quantify IgG levels binding to six ‘universal’ B-cell epitopes in COVID-19 patient sera. Analysis revealed significant variations between asymptomatic and symptomatic individuals across all six peptides of interest (**Fig. S2A**). Notably, asymptomatic patients exhibited higher IgG binding levels of the six conserved epitopes (**Figs. 4A** and **S2A**) compared to symptomatic patients, indicating a more robust humoral immune response in asymptomatic cases (**Fig. 4A**). All the six tested peptides showed significant IgG binding levels across the VOCs studied, highlighting their conserved immunogenicity (**Figs. 4A, S2A, S3A,** and **S4A**). Neutralization assays were performed to evaluate the neutralization efficiency of sera from asymptomatic and symptomatic COVID-19 patients against the different SARS-CoV-2 VOCs (**Figs. 4B, S2B, S3B,** and **S4B**). The results demonstrated notable differences in neutralization percentages between severity groups and across various VOCs. Specifically, sera from asymptomatic patients exhibited higher neutralizing antibody titers compared to sera from symptomatic patients, indicating a more potent neutralizing activity in asymptomatic cases (**Figs. 4B, S2B, S3B,** and **S4B**). In terms of age-dependent immune responses, significantly higher IgG binding levels were observed among young individuals compared to older individuals across all six peptides of interest (**Figs. 4A, S2A, S3A,** and **S4A**). Moreover, a significant increase in IgG binding levels was observed in each of the six peptides across different SARS-CoV-2 variants of concern (**Figs. 4A, S2A, S3A,** and **S4A**). Neutralization assays were conducted to assess the neutralization efficiency of young COVID-19 patient sera compared to old COVID-19 patient sera against the different SARS-CoV-2 variants of concern. The results indicated notable differences in neutralization percentages between age groups and across various VOCs, with young individuals demonstrating significantly higher neutralizing antibody titers compared to older individuals (**Figs. 4B, S2B, S3B** and **S4B**). Regarding gender-dependent immune responses, analysis of IgG binding levels revealed no significant differences between male and female patients across all six peptides of interest (**Figs. 4A, S2A, S3A,** and **S4A**). Similarly, neutralization assays showed comparable neutralization percentages between male and female patients against different VOCs (**Figs. 4B, S2B, S3B** and **S4B**).

**Figure 4.**
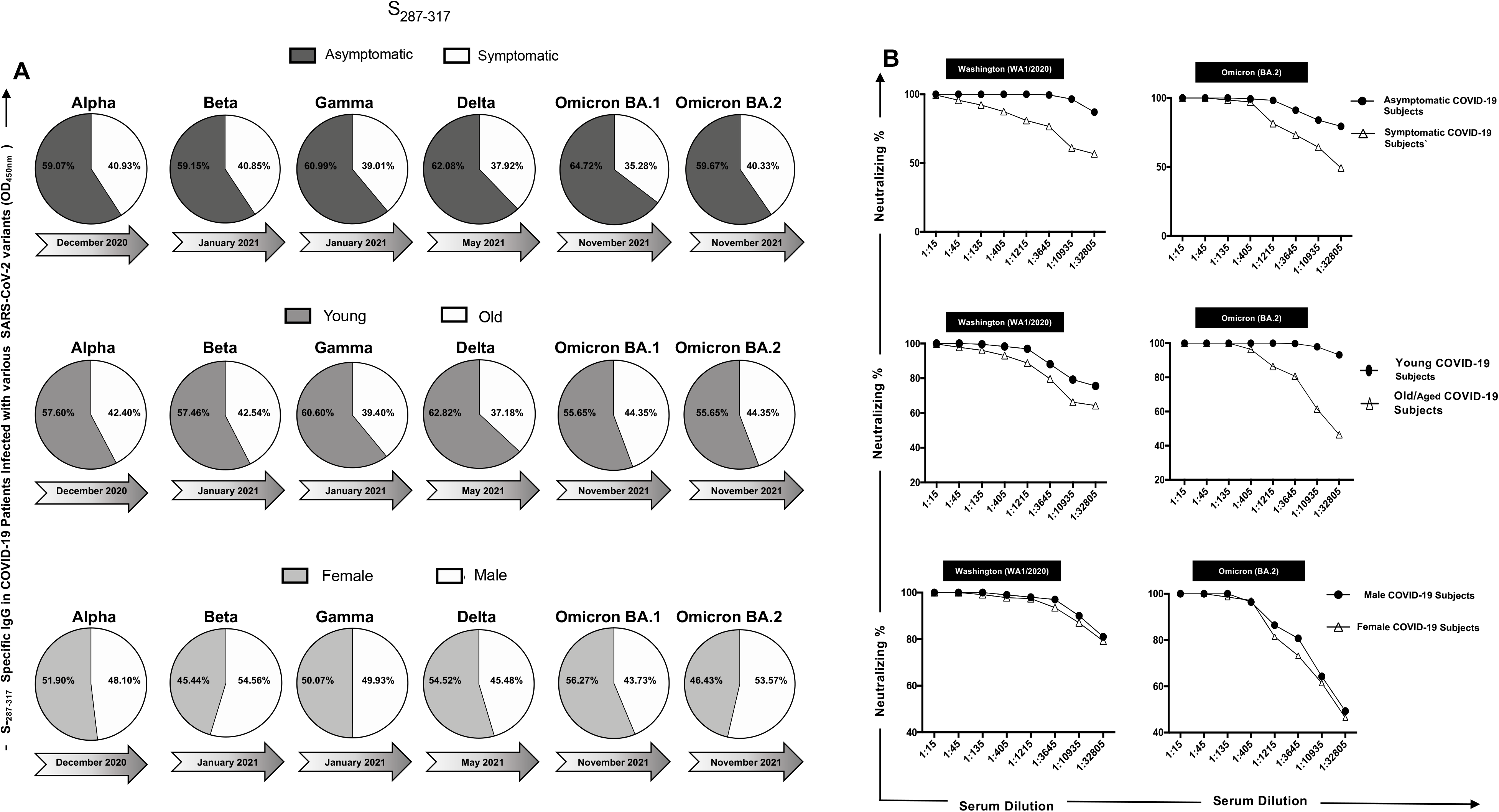
IgG levels and neutralization percentages in COVID-19 patients: a comparative analysis by ELISA and Neutralization Assay across severity, age, and gender groups: (**A**) IgG Response to Conserved Peptide in COVID-19 Patients are shown in *panel A*. The upper pie chart displays the IgG levels against a conserved peptide in symptomatic (steel) and asymptomatic (white) COVID-19 patients infected with six different SARS-CoV-2 variants of concern: Alpha, Beta, Gamma, Delta, Omicron BA.1, and Omicron BA.2. The middle pie chart shows the IgG levels against the conserved peptide in young (nickel) and old (white) COVID-19 patients infected with the same variants. The bottom pie chart represents the IgG levels against the conserved peptide in female (magnesium) and male (white) COVID-19 patients infected with the same variants. (**B**) Panel shows neutralization percent in COVID-19 patients. The upper panel illustrates the neutralization percentages in asymptomatic versus symptomatic COVID-19 patients against the Washington and Omicron BA.2 variants. The middle panel depicts the neutralization percentages in young and old COVID-19 patients against the same variants. The bottom panel displays the neutralization percentages in female and male COVID-19 patients against the same variants.

### 3. Conserved human B epitopes protect against infection and COVID-19-Like symptoms caused by SARS-CoV-2 delta variant of concern in HLA-DR0101/HLA-A0201/hACE2 triple transgenic mice

Triple transgenic HLA-A*02:01/HLA-DRB1*01:01-hACE-2 mice were intranasally vaccinated with two different AAV9-based Coronavirus vaccines: multiepitope vaccine containing 8 conserved B cell epitopes and control vaccine containing six B cell epitopes, along with a Mock vaccinated control group. ELISA and FFA assays were performed 26 days post-immunization, followed by intranasal challenge with SARS-CoV-2 Delta variant. Mice were monitored for weight loss, survival, and viral titer over 14 days post-challenge.

As illustrated in **Fig. 5B**, we observed significant protection against weight loss in the mice vaccinated with multiepitope vaccine compared to control vaccine and to the Mock vaccinated group. Additionally, a higher percentage of multiepitope vaccinated mice survived the challenge compared to the control vaccine and mock vaccinated group (**Fig. 5C**). Viral titration data demonstrated a marked decrease in viral RNA copy number in the nasopharyngeal swabs of mice vaccinated with multiepitope vaccine at days 2, 6, 10, and 14 post-challenge in comparison with control vaccine and the mock vaccinated group (**Fig. 5D**). ELISA results indicated a robust IgG binding affinity specific for the 6 “universal” B cell epitopes and the spike protein in mice vaccinated with multiepitope vaccine compared to control vaccine and mock vaccinated group (**Fig. 5E**). We also found a higher neutralization percentage in sera from mice vaccinated with multiepitope vaccine against Alpha, Beta, Epsilon, Delta, and Omicron variants compared to the control vaccine and Mock vaccinated group (**Fig. 5F**).

**Figure 5.**
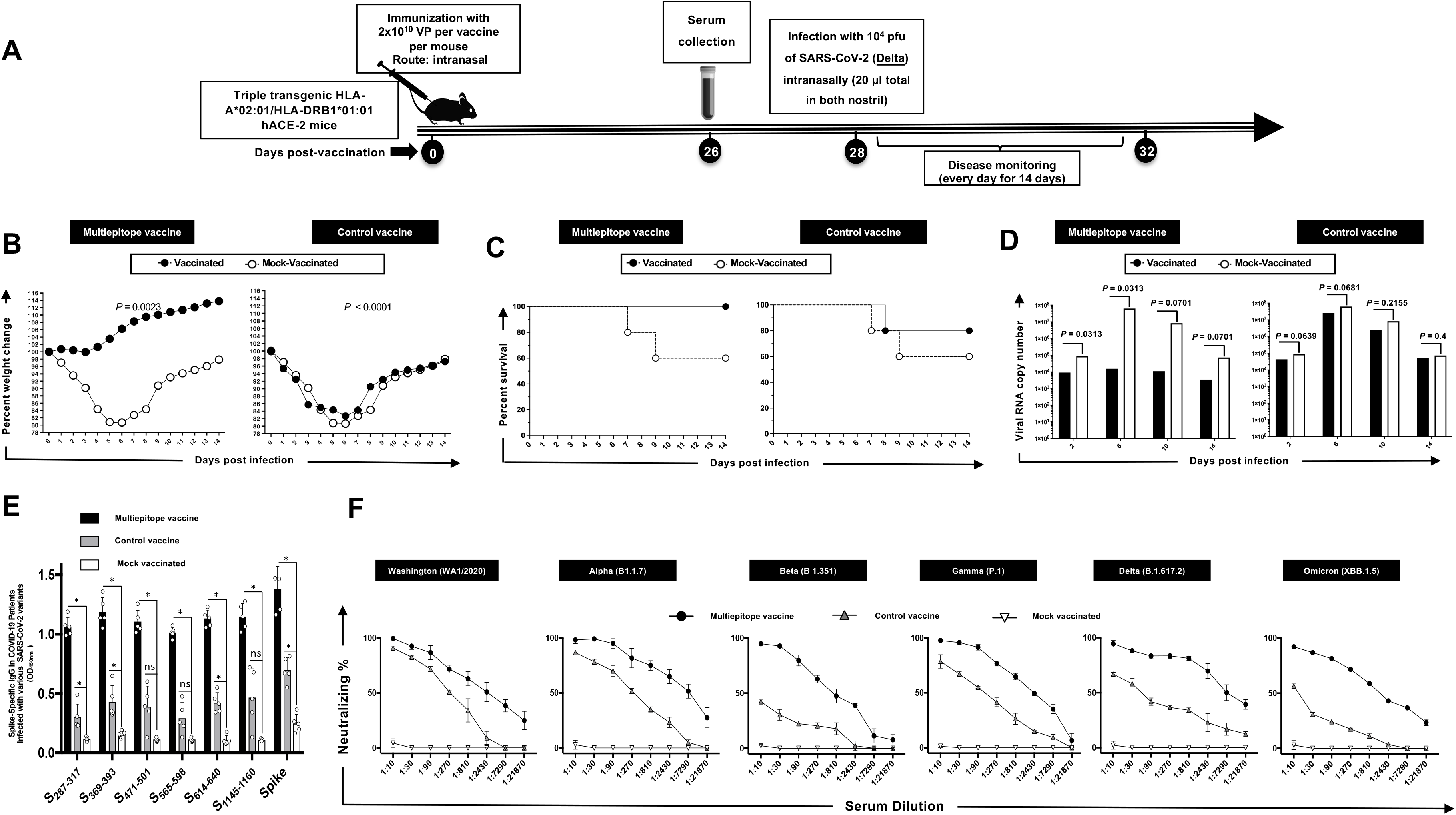
The effect of immunization with Adeno-Associated Virus 9 based multiepitope-Coronavirus vaccine incorporating conserved human B cell epitopes on COVID-19-like symptoms detected from triple transgenic HLA-A*02:01/HLA-DRB1*01:01-hACE-2 mice and infected with highly pathogenic SARS-CoV-2 Delta variant of concern: (**A**) Experimental plan to study the effect of vaccination in triple transgenic HLA-A*02:01/HLA-DRB1*01:01-hACE-2 mice. On day 0 Triple transgenic HLA-A*02:01/HLA-DRB1*01:01-hACE-2 mice (7-8-week-old, *n* = 15) were vaccinated intranasally with two different AAV9 based multiepitope vaccines 2 x 10^10^ Viral Particle per vaccine per mouse named as multiepitope vaccine containing 8 conserved B cell epitopes (*n* = 5), control vaccine containing 6 B cell epitopes (*n* = 5) and finally Mock vaccinated group that received 1XPBS (*n* = 5) were used as control. At day 26 post immunization the blood was drown for ELISA and FFA and two days later mice intranasally challenged with 20ul of SARS-CoV-2 Delta (B.1.617.2) variant of concern at 1 x 10^4^ pfu. Mice were followed for wight loss, survival, and viral titer for 14 days. (**B**) Data showing average percent weight change each day post immunization to the body weight on the day of infection. (**C**) shows the percentage survival detected in mice groups that received either multiepitope vaccine or control vaccine and finally the mock vaccinated group. (**D**) Viral titration data showing viral RNA copy number in the nasopharyngeal swabs of each group at days 2, 6, 10 and 14 post challenge Delta (B.1.617.2) variant. The IgG binding affinity specific for 6 “universal” B cell epitopes as well as the spike protein measured by ELISA are shown in *panel* (**E**). The (**F**) *panel* represents neutralization percent by sera from mice that were given multiepitope vaccine, control vaccine or Mock vaccinated group against Alpha (B.1.1.7), Beta (B.1.351), Epsilon (B.1.427/B.1.429), Delta (B.1.617.2), and Omicron (XBB1.5). Bars represent means ± SEM. *P* values are calculated using unpaired *t*-test, comparing results obtained in vaccinated vs. mock-vaccinated mice.

## DISCUSSION

Since the emergence of SARS-CoV-2 in late 2019, the identification and understanding of B cell epitopes have become pivotal in the development of effective vaccines and therapeutic strategies against COVID-19. B cell epitopes, which are specific regions on antigens recognized by antibodies, play a crucial role in mediating the neutralization of the virus. The continuous emergence of SARS-CoV-2 variants, including the recent Omicron sub-variants, has complicated efforts to control the COVID-19 pandemic. Despite the significant impact of Spike-based COVID-19 vaccines, their effectiveness against newer variants is waning (19, 20), underscoring the pressing need for a next-generation SARS-CoV-2 vaccine that contains a repertoire of conserved B cell epitopes. The exploration of conserved B cell epitopes in a universal coronavirus vaccine offers a promising path to enhancing immunity against the multitude of SARS-CoV-2 variants and sub-variants, thereby potentially attenuating the persistent threat posed by the pandemic.

B cell epitopes are short sequences or structures on an antigen recognized by antibodies produced by B cells. Neutralization occurs when these antibodies bind to viral epitopes and block the virus from entering host cells. For SARS-CoV-2, the primary target of neutralizing antibodies is the spike (S) protein, which facilitates viral entry into human cells by binding to the ACE2 receptor (21). The spike protein of SARS-CoV-2 is composed of two subunits: S1 and S2. The S1 subunit includes the receptor-binding domain (RBD), which is a major B cell epitope. Antibodies targeting the RBD are highly effective at neutralizing the virus because they prevent the spike protein from interacting with the ACE2 receptor (22). Other important epitopes include the N-terminal domain (NTD) of the S1 subunit and the S2 subunit, although the RBD remains the primary focus for neutralizing responses (23–25). However, it was noticed that the emerging variants of SARS-CoV-2 plays a role in B cell epitope recognition. Since 2019, SARS-CoV-2 has undergone several mutations, leading to the emergence of new variants with changes in the spike protein. These variants, including Alpha, Beta, Delta, and Omicron, have introduced mutations in the RBD and other regions, potentially altering the B cell epitopes (26). For instance, the Omicron variant has multiple mutations in the RBD, which can reduce the effectiveness of antibodies generated by previous infection or vaccination (27). Understanding these changes is crucial for maintaining vaccine efficacy and developing new therapeutic strategies.

The design of COVID-19 vaccines has been heavily influenced by the need to elicit a strong antibody response against key B cell epitopes. mRNA vaccines, such as those developed by Pfizer-BioNTech and Moderna, encode the spike protein and stimulate an immune response targeting the RBD (28–30). Viral vector vaccines, like AstraZeneca’s, also focus on the spike protein but use a different delivery mechanism. The effectiveness of these vaccines in neutralizing SARS-CoV-2 largely depends on the ability of the induced antibodies to recognize and bind to critical epitopes on the spike protein (31). Several studies regarding the effectiveness of the current modified messenger RNA (mRNA) vaccines indicate that these vaccines had reduced levels of neutralizing antibodies against recent SARS-CoV-2 variants compared to earlier variants (19, 32). In this report, we have identified 6 conserved B-cell epitopes among all known SARS-CoV-2 variants, previous SARS and MERS coronavirus strains, and strains specific to different species that were reported to be hosts for SARS/MERS (bat, civet cat, pangolin, and camel). In this study, we used a combination of these highly conserved B cell epitopes and incorporated them in a multi-epitope pan-variant SARS-CoV-2 vaccine that contain CD8^+^ and CD4^+^ T- and B- cell epitopes.

We demonstrated that among the seventeen conserved B-cell epitopes, six (S_287-317_, S_369-393_, S_471-501_, S_565-598_, S_614-640_, S_1133-1160_) were highly immunogenic. The magnitude of IgG response consistently of these six peptides exhibit high immunogenicity and stable responses across different SARS-CoV-2 VOCs, when compared to other conserved peptides. Suggesting that these peptides elicit stronger and more consistent IgG responses over time, despite the emergence of diverse viral variants compared to their counterparts, providing a clear indication of their reliability and effectiveness in eliciting immune responses. These epitopes exhibited robust immunogenicity and demonstrated conservation across multiple SARS-CoV-2 variants. This analysis revealed that over time, the immunogenicity of conserved epitopes remained stable and higher compared to non-conserved epitopes. Conversely, the immunogenicity of non-conserved epitopes exhibited a declining trend over time. Upon selecting six conserved epitopes (S_287-317_, S_369-393_, S_471-501_, S_565-598_, S_614-640_, S_1133-1160_) as highly immunogenic peptides, we subsequently placed additional focus on these peptides and correlated their IgG level with severity, age, and gender. Notably, asymptomatic patients showed a significantly higher IgG binding levels to the six conserved epitopes (S_287-317_, S_369-393_, S_471-501_, S_565-598_, S_614-640_, S_1133-1160_) across different SARS-CoV-2 VOCs Alpha (B.1.1.7), Beta (B.1.351), Gamma (P.1), Delta (B.1.617.2), Omicron (BA.1) and Omicron (BA.2) when compared to symptomatic patients. Furthermore, when investigating the neutralization levels using sera from asymptomatic and symptomatic individuals across various VOCs, we also demonstrated that sera from asymptomatic patients exhibited higher neutralizing antibody titers compared to sera from symptomatic patients, indicating a more potent neutralizing activity in asymptomatic. Our studies suggest a potential correlation between elevated IgG levels specific to certain peptides and increased neutralization activity in asymptomatic individuals, pointing towards a more robust immune response in this cohort. Altogether, these findings imply that the magnitude of the peptide-specific IgG response may influence the ability to neutralize the virus, highlighting the importance of further exploring this relationship in understanding COVID-19 pathogenesis and immune response dynamics.

This study investigated age-related immune responses and found a trend of significantly higher IgG binding levels among younger individuals in comparison to their older counterparts across all six peptides examined suggesting a potential age-dependent variation in the humoral immune response to SARS-CoV-2 infection, with younger individuals exhibiting a more robust antibody response to the viral epitopes of interest. Furthermore, our analysis revealed a consistent increase in IgG binding levels across all six peptides when considering different VOCs of SARS-CoV-2. This observation indicates that the epitopes targeted by these antibodies remain immunogenic and conserved across various viral strains, underscoring their potential significance as vaccine targets against evolving viral variants. To further assess the functional relevance of these age-related differences in antibody responses, we conducted neutralization assays using sera from young and old COVID-19 patients against different VOCs of SARS-CoV-2. Notable increases were observed in neutralization efficiency between age groups and across various viral variants. Specifically, young individuals exhibited significantly higher neutralizing antibody titers compared to their older counterparts, indicating a potential age-associated variation in the ability to neutralize the virus. These findings suggest that age may play a critical role in shaping the magnitude and efficacy of the humoral immune response to SARS-CoV-2 infection. Understanding these age-related differences in immune responses could have important implications for vaccine design and prioritization, as well as for informing strategies aimed at enhancing vaccine efficacy across different age demographics. Further research is warranted to elucidate the underlying mechanisms driving these age-dependent variations in immune responses and to assess their implications for COVID-19 disease severity and vaccine effectiveness. In examining gender-dependent immune responses, our analysis of IgG binding levels across the six peptides of interest revealed no significant differences between male and female patients. Similarly, we observed comparable neutralization percentages between male and female patients across different VOCs when performing neutralization assays to evaluate the efficacy of neutralizing antibodies, This study further suggests that gender may not be a significant factor influencing the humoral immune response to SARS-CoV-2 infection, at least in terms of IgG binding and neutralizing antibody titers.

After identifying six highly conserved epitopes (S_287-317_, S_369-393_, S_471-501_, S_565-598_, S_614-640_, S_1133-1160_), we evaluated the in vivo efficacy of a multiepitope vaccine incorporating these highly conserved B cell epitopes along with CD4^+^ T cell and CD8^+^ T cell epitopes using a triple transgenic HLA-A*02:01/HLA-DRB1*01:01-hACE-2 mouse model. Our findings demonstrated significant protective effects of the multiepitope vaccine against SARS-CoV-2 (Delta variant) challenge. Mice immunized with the multiepitope vaccine showed substantial protection against weight loss and death. The weight loss and survival found herein agree with previous reports in the context to Delta (B.1.617.2) (33). Viral titration analysis revealed a significant reduction in viral RNA copy numbers in the nasopharyngeal swabs of mice vaccinated with the multiepitope vaccine at various time points post-challenge (days 2, 6, 10, and 14). This indicates effective viral clearance in vaccinated mice compared to the control groups. Furthermore, ELISA results demonstrated robust IgG binding affinity specific to the six “universal” B cell epitopes and the spike protein in mice vaccinated with the multiepitope vaccine. Importantly, we observed a higher neutralization percentage in sera from mice vaccinated with the multiepitope vaccine against various SARS-CoV-2 variants, including Alpha, Beta, Epsilon, Delta, and Omicron, compared to the control groups. This highlights the vaccine’s ability to induce a strong humoral immune response targeting conserved epitopes underscoring the potential of the multiepitope vaccine to confer broad protection against emerging VOCs. Overall, our results demonstrate the promising efficacy of the multiepitope vaccine in providing protection against SARS-CoV-2 infection and highlight its potential as a candidate for further preclinical and clinical development.

There are several studies on epitope profiling in existing COVID-19 mRNA vaccines (34, 35). One study mapped immunogenic amino acid motifs and linear epitopes of the primary sequence of the SARS-CoV-2 spike protein that induce IgG in recipients of the Pfizer-BioNTech COVID-19 mRNA vaccine (34). The data revealed various distinctive amino acid motifs recognized by vaccine-elicited IgG, some of which mimic three-dimensional conformation (mimotopes) and are identical to dominant linear epitopes in the C-terminal region of the spike protein observed in SARS-CoV, bat coronaviruses, and epitopes triggering IgG during natural infection. However, these epitopes have limited homology to the spike protein of non-pathogenic human coronaviruses (34). Another study highlighted high-resolution linear epitope profiling of Pfizer-BioNTech COVID-19 mRNA vaccine recipients and COVID-19 patients showed that vaccine-induced antibodies targeting the viral spike receptor-binding domain (RBD) have a broader distribution across the RBD compared to antibodies induced by natural infection (35). Furthermore, mutation panel assays targeting viral variants of concern demonstrated that the epitope repertoire induced by the mRNA vaccine is rich in breadth, potentially conferring resistance against viral evolutionary escapes in the future. This represents a significant advantage of vaccine-induced immunity (35). The identified epitopes in COVID-19 mRNA vaccines may serve as a basis for further research on immune escape, viral variants, and the design of vaccines and therapies.

SARS-CoV-2 shares sequence, structural, and functional homology with SARS-CoV and MERS-CoV (36). Antibodies against SARS-CoV spike protein can inhibit SARS-CoV-2 binding to ACE2 (37). The spike protein, vital for receptor binding, possesses conserved sequences and high immunogenicity, making it a priority in COVID-19 vaccine development. Out of the current clinical vaccine candidates, 32% are recombinant protein vaccines, while RNA vaccines, viral vector-based vaccines, and inactivated virus vaccines account for 23%, 13%, and 13% respectively (38, 39). Many of these candidates use either full-length spike protein or only parts from the spike protein various lengths of the receptor-binding domain (RBD) to induce potent neutralizing antibody responses. However, it is important to note that mutations in spike protein can impact the effectiveness of those COVID-19 vaccines based only on the spike protein (36). Ongoing research aims to identify additional B cell epitopes and understand their role in neutralization, especially in the context of evolving variants. This includes studying the cross-reactivity of antibodies and the development of broadly neutralizing antibodies that can target conserved regions of the spike protein across different variants (40, 41). Advances in structural biology and immunology will continue to enhance our understanding of B cell epitopes and their role in combating SARS-CoV-2. Understanding the role of B cell epitopes in neutralizing SARS-CoV-2 is crucial for ongoing efforts to control the pandemic. Continued research into these epitopes will be vital for adapting vaccines and treatments to effectively address emerging variants and ensure robust protection against COVID-19.

The threat of COVID-19 still remains serious, with persistently high rates of illness and mortality worldwide. Our findings underscore the efficacy and potential of a multiepitope vaccine designed to target conserved B-cell epitopes as well as CD4^+^ and CD8^+^ T cell epitopes. Our strategy of incorporating selected highly conserved B cell epitopes into COVID-19 vaccines presents a pivotal approach in the fight against emerging variants. By putting to use the power of both humoral and cellular immunity, a multiepitope vaccines hold significant promise and provides a broader and more durable protection against multiple SARS-CoV-2 variants. This investigation highlights the importance of incorporating B-cell epitopes into next-generation vaccine strategies for enhanced protection against evolving viral threats.

**Table 2.** Screening COVID-19 patients involved assessing SARS-CoV-2 variants, age, gender, and severity: Screening process of COVID-19 patients (*n* = 210) into Asymptomatic (*n* = 113) and Symptomatic (*n* = 85) categories based on clinical parameters. the groups were segregated based on gender into females (*n* = 96) and males (*n* = 102), as well as based on age into young (age between 19 and 49 years old) individuals (*n* = 111) and old (age between 50 and 89 years old) individuals (*n* = 87). Blood and nasopharyngeal swabs were collected from all the subjects and a qRT-PCR assay was performed. Six novel nonsynonymous mutations (Δ69-70, Δ242-244, N501Y, E484K, L452R, and T478K) were used to identify the haplotypes unique to different SARS-CoV-2 variants of concern (Omicron (B.1.1.529 (BA.1)), Omicron (B.1.1.529 (BA.2)), Alpha (B.1.1.7), Beta (B.1.351), Gamma (P.1), Delta (B.1.617.2), and Epsilon (B.1.427/B.1.429)).

|  | Symptom |  | Gender |  | Age |  |  |
| --- | --- | --- | --- | --- | --- | --- | --- |
| Variants of Concern | SYMP (%) | ASYMP (%) | Male (%) | Female (%) | Young (%) | Old (%) | Total Sample Size (%) |
| B.1.1.7 (Alpha) | 10 (45.4) | 12 (54.6) | 12 (54.5) | 10 (45.5) | 14 (63.6) | 8 (36.4) | 22 (11.1) |
| B.1.351( Beta) | 8 (47.1) | 9 (52.9) | 8 (47.1) | 9 (52.9) | 11 (64.7) | 6 (35.3) | 17 (8.6) |
| P.1. (Gamma) | 6 (31.5) | 13 (68.5) | 9 (47.3) | 10 (52.7) | 10 (52.7) | 9 (47.3) | 19 (9.6) |
| B.1.617.2 (Delta) | 18 (40.9) | 26 (59.1) | 24 (54.5) | 20 (45.5) | 24 (54.5) | 20 (45.5) | 44 (22.2) |
| B.1.427/B.1.429 (Epsilon) | 16 (51.6) | 15 (48.4) | 17 (54.8) | 14 (45.2) | 20 (64.5) | 11 (35.5) | 31 (15.7) |
| B.1.1.529 (BA.1) Omicron | 17 (48.5) | 18 (51.5) | 16 (45.7) | 19 (54.3) | 16 (45.7) | 19 (54.3) | 35 (17.7) |
| B.1.1.529 (BA.2) Omicron | 10 (33.3) | 20 (66.7) | 16 (53.3) | 14 (46.7) | 16 (53.3) | 14 (46.6) | 30 (15.1) |

The first-generation Spike-alone-based COVID-19 vaccines have successfully reduced the risk of hospitalization, serious illness, and death caused by SARS-CoV-2 infections (1–3, 11, 12). However, waning immunity induced by these vaccines failed to prevent immune escape by many VOCs that have emerged from 2020 to 2024, resulting in a prolonged COVID-19 pandemic.

## ACKNOWLEDGMENTS

This work is supported by the Fast-Grant PR12501 from Emergent Ventures, by a Gavin Herbert Eye Institute internal grant, by Public Health Service Research grants AI158060, AI174383, AI150091, AI143348, AI147499, AI143326, AI138764, AI124911, and AI110902 from the National Institutes of Allergy and Infectious Diseases (NIAID) to LBM.

## REFERENCES

1. Prakash S, Srivastava R, Coulon PG, Dhanushkodi NR, Chentoufi AA, Tifrea DF, et al. Genome-Wide B Cell, CD4(+), and CD8(+) T Cell Epitopes That Are Highly Conserved between Human and Animal Coronaviruses, Identified from SARS-CoV-2 as Targets for Preemptive Pan-Coronavirus Vaccines. J Immunol. 2021;206(11):2566–82.

2. Prakash S, Dhanushkodi NR, Zayou L, Ibraim IC, Quadiri A, Coulon PG, et al. Cross-Protection Induced by Highly Conserved Human B, CD4 (+,) and CD8 (+) T Cell Epitopes-Based Coronavirus Vaccine Against Severe Infection, Disease, and Death Caused by Multiple SARS-CoV-2 Variants of Concern. Frontiers In Immunololgy. 2024;29(12):45–56.

3. Pedersen SF, Ho YC. SARS-CoV-2: A Storm is Raging. J Clin Invest. 2020.

4. Bellocchi MC, Scutari R, Carioti L, Iannetta M, Marchegiani G, Piermatteo L, et al. Frequency of Atypical Mutations in the Spike Glycoprotein in SARS-CoV-2 Circulating from July 2020 to July 2022 in Central Italy: A Refined Analysis by Next Generation Sequencing. Viruses. 2023;15(8).

5. Desmecht S, Tashkeev A, El Moussaoui M, Marechal N, Peree H, Tokunaga Y, et al. Kinetics and Persistence of the Cellular and Humoral Immune Responses to BNT162b2 mRNA Vaccine in SARS-CoV-2-Naive and -Experienced Subjects: Impact of Booster Dose and Breakthrough Infections. Front Immunol. 2022;13:863554.

6. Sharma S, Vercruysse T, Sanchez-Felipe L, Kerstens W, Rasulova M, Bervoets L, et al. Updated vaccine protects against SARS-CoV-2 variants including Omicron (B.1.1.529) and prevents transmission in hamsters. Nat Commun. 2022;13(1):6644.

7. Washington NL, Gangavarapu K, Zeller M, Bolze A, Cirulli ET, Schiabor Barrett KM, et al. Emergence and rapid transmission of SARS-CoV-2 B.1.1.7 in the United States. Cell. 2021;184(10):2587-94 e7.

8. Liu H, Iketani S, Zask A, Khanizeman N, Bednarova E, Forouhar F, et al. Development of optimized drug-like small molecule inhibitors of the SARS-CoV-2 3CL protease for treatment of COVID-19. Nat Commun. 2022;13(1):1891.

9. Konings F, Perkins MD, Kuhn JH, Pallen MJ, Alm EJ, Archer BN, et al. SARS-CoV-2 Variants of Interest and Concern naming scheme conducive for global discourse. Nat Microbiol. 2021;6(7):821–3.

10. Kumar S, Thambiraja TS, Karuppanan K, Subramaniam G. Omicron and Delta variant of SARS-CoV-2: A comparative computational study of spike protein. J Med Virol. 2022;94(4):1641–9.

11. Park T, Hwang H, Moon S, Kang SG, Song S, Kim YH, et al. Vaccines against SARS-CoV-2 variants and future pandemics. Expert Rev Vaccines. 2022;21(10):1363–76.

12. Evans JP, Liu SL. Challenges and Prospects in Developing Future SARS-CoV-2 Vaccines: Overcoming Original Antigenic Sin and Inducing Broadly Neutralizing Antibodies. J Immunol. 2023;211(10):1459–67.

13. Lyke KE, Atmar RL, Islas CD, Posavad CM, Szydlo D, Paul Chourdhury R, et al. Rapid decline in vaccine-boosted neutralizing antibodies against SARS-CoV-2 Omicron variant. Cell Rep Med. 2022;3(7):100679.

14. Shi J, Zheng J, Zhang X, Tai W, Compas R, Deno J, et al. A T cell-based SARS-CoV-2 spike protein vaccine provides protection without antibodies. JCI Insight. 2024;9(5).

15. Gatz SA, Pohla H, Schendel DJ. A PCR-SSP method to specifically select HLA-A*0201 individuals for immunotherapeutic studies. Tissue antigens. 2000;55(6):532–47.

16. Olerup O, Zetterquist H. HLA-DRB1*01 subtyping by allele-specific PCR amplification: a sensitive, specific and rapid technique. Tissue antigens. 1991;37(5):197–204.

17. Buchfink B, Reuter K, Drost HG. Sensitive protein alignments at tree-of-life scale using DIAMOND. Nat Methods. 2021;18(4):366–8.

18. Wang H, Jean S, Eltringham R, Madison J, Snyder P, Tu H, et al. Mutation-Specific SARS-CoV-2 PCR Screen: Rapid and Accurate Detection of Variants of Concern and the Identification of a Newly Emerging Variant with Spike L452R Mutation. J Clin Microbiol. 2021;59(8):e0092621.

19. Duchene S, Featherstone L, Haritopoulou-Sinanidou M, Rambaut A, Lemey P, Baele G. Temporal signal and the phylodynamic threshold of SARS-CoV-2. Virus Evol. 2020;6(2):veaa061.

20. Rubin R. COVID-19 Vaccines vs Variants-Determining How Much Immunity Is Enough. JAMA. 2021;325(13):1241–3.

21. Hoffmann M, Kleine-Weber H, Schroeder S, Kruger N, Herrler T, Erichsen S, et al. SARS-CoV-2 Cell Entry Depends on ACE2 and TMPRSS2 and Is Blocked by a Clinically Proven Protease Inhibitor. Cell. 2020;181(2):271–80 e8.

22. Barton MI, MacGowan SA, Kutuzov MA, Dushek O, Barton GJ, van der Merwe PA. Effects of common mutations in the SARS-CoV-2 Spike RBD and its ligand, the human ACE2 receptor on binding affinity and kinetics. Elife. 2021;10.

23. Liu L, Iketani S, Guo Y, Chan JF, Wang M, Liu L, et al. Striking antibody evasion manifested by the Omicron variant of SARS-CoV-2. Nature. 2022;602(7898):676-81.

24. Liu L, Wang P, Nair MS, Yu J, Rapp M, Wang Q, et al. Potent neutralizing antibodies against multiple epitopes on SARS-CoV-2 spike. Nature. 2020;584(7821):450-6.

25. Carabelli AM, Peacock TP, Thorne LG, Harvey WT, Hughes J, Consortium C-GU, et al. SARS-CoV-2 variant biology: immune escape, transmission and fitness. Nat Rev Microbiol. 2023;21(3):162–77.

26. Planas D, Veyer D, Baidaliuk A, Staropoli I, Guivel-Benhassine F, Rajah MM, et al. Reduced sensitivity of SARS-CoV-2 variant Delta to antibody neutralization. Nature. 2021;596(7871):276-80.

27. Chen RE, Zhang X, Case JB, Winkler ES, Liu Y, VanBlargan LA, et al. Resistance of SARS-CoV-2 variants to neutralization by monoclonal and serum-derived polyclonal antibodies. Nat Med. 2021;27(4):717–26.

28. Cele S, Gazy I, Jackson L, Hwa SH, Tegally H, Lustig G, et al. Escape of SARS-CoV-2 501Y.V2 from neutralization by convalescent plasma. Nature. 2021;593(7857):142–6.

29. El Sahly HM, Baden LR, Essink B, Doblecki-Lewis S, Martin JM, Anderson EJ, et al. Efficacy of the mRNA-1273 SARS-CoV-2 Vaccine at Completion of Blinded Phase. The New England journal of medicine. 2021;385(19):1774–85.

30. Polack FP, Thomas SJ, Kitchin N, Absalon J, Gurtman A, Lockhart S, et al. Safety and Efficacy of the BNT162b2 mRNA Covid-19 Vaccine. The New England journal of medicine. 2020;383(27):2603–15.

31. Slaoui M, Hepburn M. Developing Safe and Effective Covid Vaccines - Operation Warp Speed’s Strategy and Approach. N Engl J Med. 2020;383(18):1701–3.

32. Hao L, Hsiang TY, Dalmat RR, Ireton R, Morton JF, Stokes C, et al. Dynamics of SARS-CoV-2 VOC Neutralization and Novel mAb Reveal Protection against Omicron. Viruses. 2023;15(2).

33. Prakash S, Dhanushkodi NR, Zayou L, Ibraim IC, Quadiri A, Coulon PG, et al. Cross-protection induced by highly conserved human B, CD4(+), and CD8(+) T-cell epitopes-based vaccine against severe infection, disease, and death caused by multiple SARS-CoV-2 variants of concern. Front Immunol. 2024;15:1328905.

34. Amanat F, Thapa M, Lei T, Ahmed SMS, Adelsberg DC, Carreño JM, et al. SARS-CoV-2 mRNA vaccination induces functionally diverse antibodies to NTD, RBD, and S2. Cell. 2021;184(15):3936–48.e10.

35. Nitahara Y, Nakagama Y, Kaku N, Candray K, Michimuko Y, Tshibangu-Kabamba E, et al. High-Resolution Linear Epitope Mapping of the Receptor Binding Domain of SARS-CoV-2 Spike Protein in COVID-19 mRNA Vaccine Recipients. Microbiol Spectr. 2021;9(3):e0096521.

36. Wu A, Peng Y, Huang B, Ding X, Wang X, Niu P, et al. Genome Composition and Divergence of the Novel Coronavirus (2019-nCoV) Originating in China. Cell Host Microbe. 2020;27(3):325–8.

37. Salvatori G, Luberto L, Maffei M, Aurisicchio L, Roscilli G, Palombo F, Marra E. SARS-CoV-2 SPIKE PROTEIN: an optimal immunological target for vaccines. J Transl Med. 2020;18(1):222.

38. Liang JG, Su D, Song TZ, Zeng Y, Huang W, Wu J, et al. S-Trimer, a COVID-19 subunit vaccine candidate, induces protective immunity in nonhuman primates. Nat Commun. 2021;12(1):1346.

39. Nelde A, Bilich T, Heitmann JS, Maringer Y, Salih HR, Roerden M, et al. SARS-CoV-2-derived peptides define heterologous and COVID-19-induced T cell recognition. Nat Immunol. 2020.

40. Li Y, Xu Z, Lei Q, Lai DY, Hou H, Jiang HW, et al. Antibody landscape against SARS-CoV-2 reveals significant differences between non-structural/accessory and structural proteins. Cell Rep. 2021;36(2):109391.

41. Qi H, Liu B, Wang X, Zhang L. The humoral response and antibodies against SARS-CoV-2 infection. Nat Immunol. 2022;23(7):1008–20.

