## Supplemental Figures for "Dynamics of Spike-Specific Neutralizing Antibodies Across Five-Year Emerging SARS-CoV-2 Variants of Concern Reveal Conserved Epitopes that Protect Against Severe COVID-19"

*Supplemental Fig. S1*

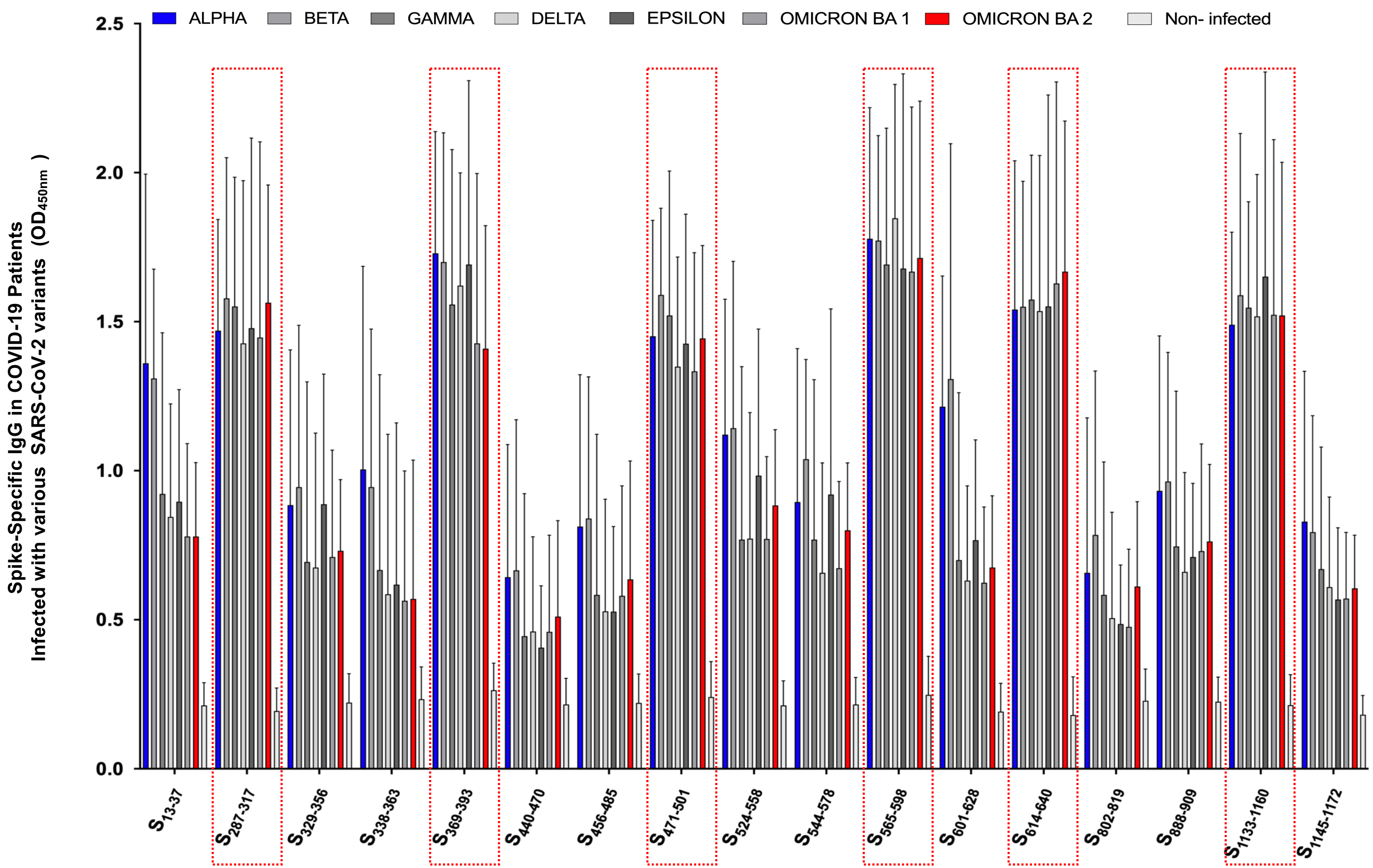

**Supplemental Figure S1: Evaluation of IgG binding to conserved B cell "asymptomatic" epitopes.** Graph shows the optical density for anti SARS-CoV-2 peptide specific IgG measured in sera from different groups of COVID-19 infected with highly pathogenic SARS-CoV-2 variants of concern. Dotted lines indicate selected peptides demonstrating notably high immunogenicity among the 17 peptides analyzed. The bottom panel identifies sequence homology analysis degree of the conservancy of the immunodominant B cell epitopes among SARS-CoV-2 variants of concern.

*Supplemental Fig. S2*

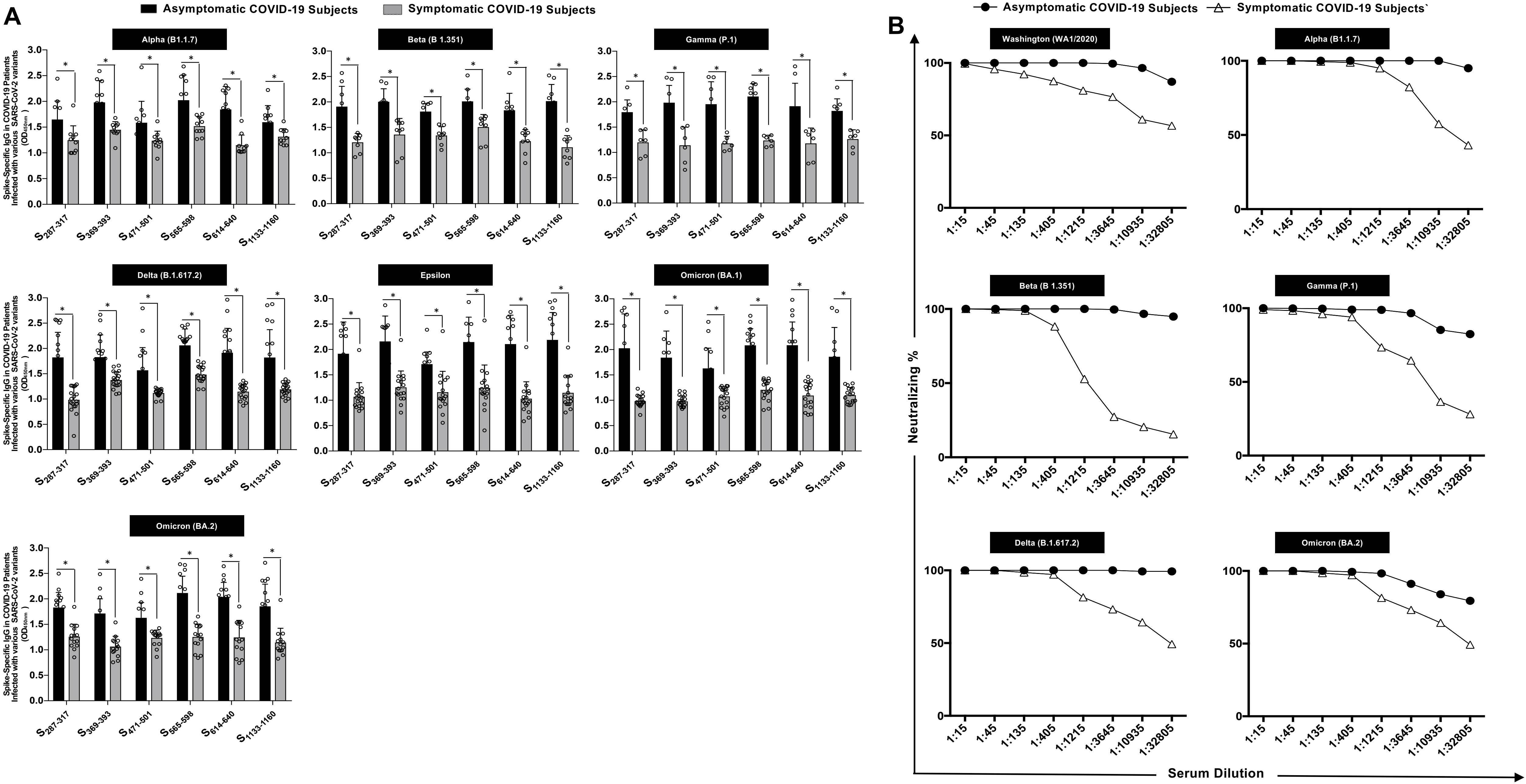

**Supplemental Figure S2. Severity-dependent immune responses against 'universal' B-cell epitopes in COVID-19 patients exposed to different SARS-CoV-2 variants of concern: Highly conserved COVID-19 peptide immunogenicity evaluation.** Bar graphs show the peptide binding IgG level for the 6 “universal” B cell epitopes meseaured by ELISA as (*panel* **A**). Serum samples were obtained from COVID-19 patients infected with various SARS-CoV-2 variants of concern (VOC) including Alpha (B.1.1.7), Beta (B.1.351), Epsilon (B.1.427/B.1.429), Delta (B.1.617.2), and Omicron (BA.1 and BA.2), and segregated into two distinct groups based on severity level, categorized as "Asymptomatic" and "Symptomatic". Neutralization (%) by sera from these patients against the different VOCs of SARS-CoV-2: Alpha (B.1.1.7), Beta (B.1.351), Epsilon (B.1.427/B.1.429), Delta (B.1.617.2), and Omicron (BA.2) is presented in (*panel* **B**)**.** Bars represent means ± SEM. Data were analyzed by student's *t*-test and multiple t-tests. Results were considered statistically significant at $P < 0.05$. Statistical correction for multiple comparisons was applied using the Holm-Sidak method.

*Supplemental Fig. S3*

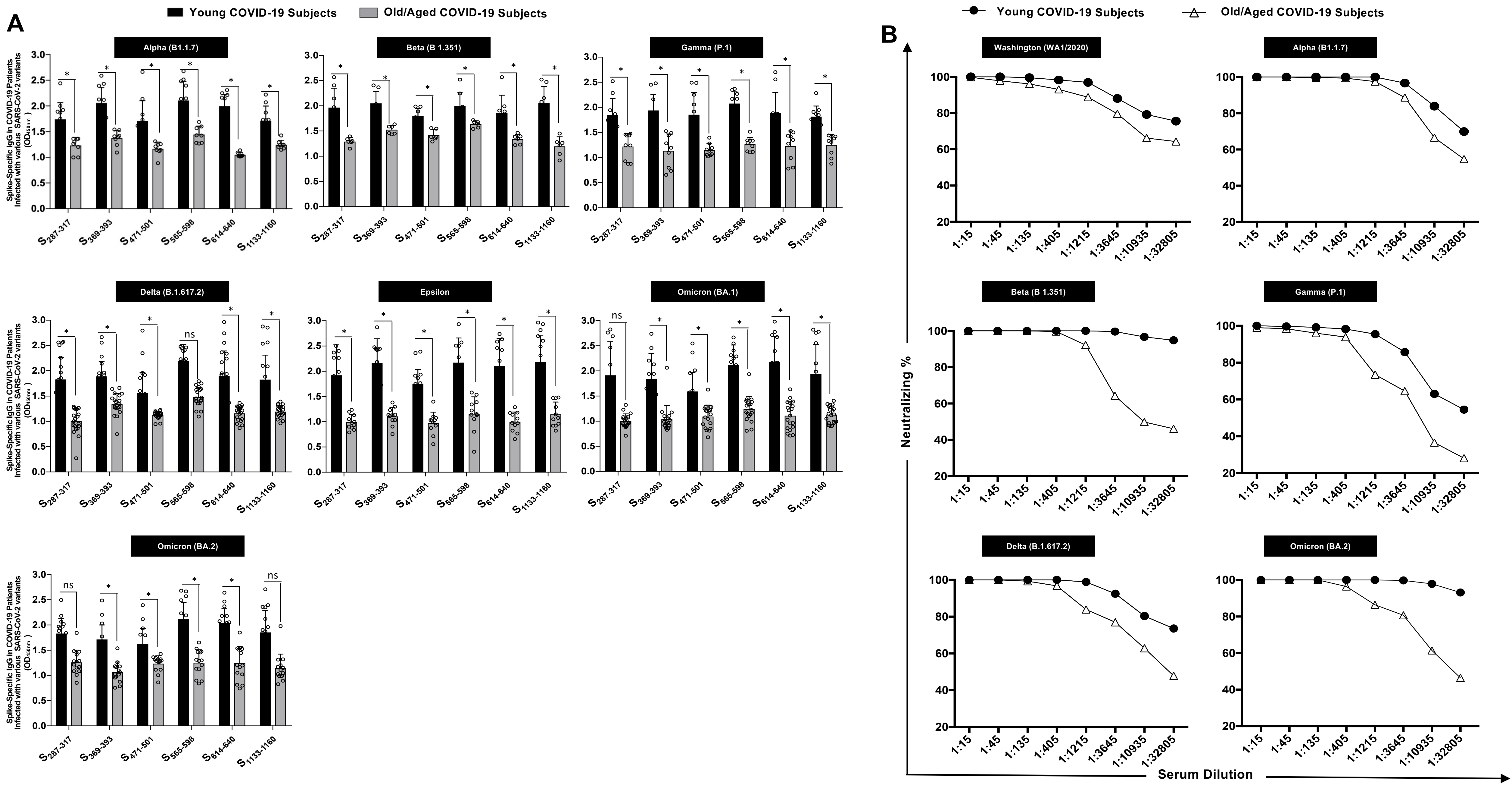

**<u>Supplemental Figure S3</u>. Age-dependent immune responses against 'universal' B-cell epitopes in COVID-19 patients exposed to different SARS-CoV-2 variants of concern: Highly conserved COVID-19 peptide immunogenicity evaluation.** Bar graphs represent the peptide binding IgG level for the 6 “universal” B cell epitopes meseaured by ELISA as shown in *panel* (**A**). Serum was obtained from COVID-19 patients that were infected with one of the following six SARS-CoV-2 different variants of concern (VOC) Alpha (B.1.1.7), Beta (B.1.351), Epsilon (B.1.427/B.1.429), Delta (B.1.617.2), and Omicron (BA.1) and Omicron (BA.2) and then segregated into two distinct age groups, categorized as "Old" and "Young". The (**B**) *panel* represents Neutralization (%) by sera from COVID-19 patients that were infected with one of the six different variants of concern (VOC) of SARS-CoV-2 and then segregated into two distinct age groups; "Old" and "Young" against Alpha (B.1.1.7), Beta (B.1.351), Epsilon (B.1.427/B.1.429), Delta (B.1.617.2), and Omicron (BA.2)**.** Bars represent means ± SEM. Data were analyzed by student's *t*-test and multiple t-tests. Results were considered statistically significant at $P < 0.05$. Statistical correction for multiple comparisons was applied using the Holm-Sidak method.

*Supplemental Fig. S4*

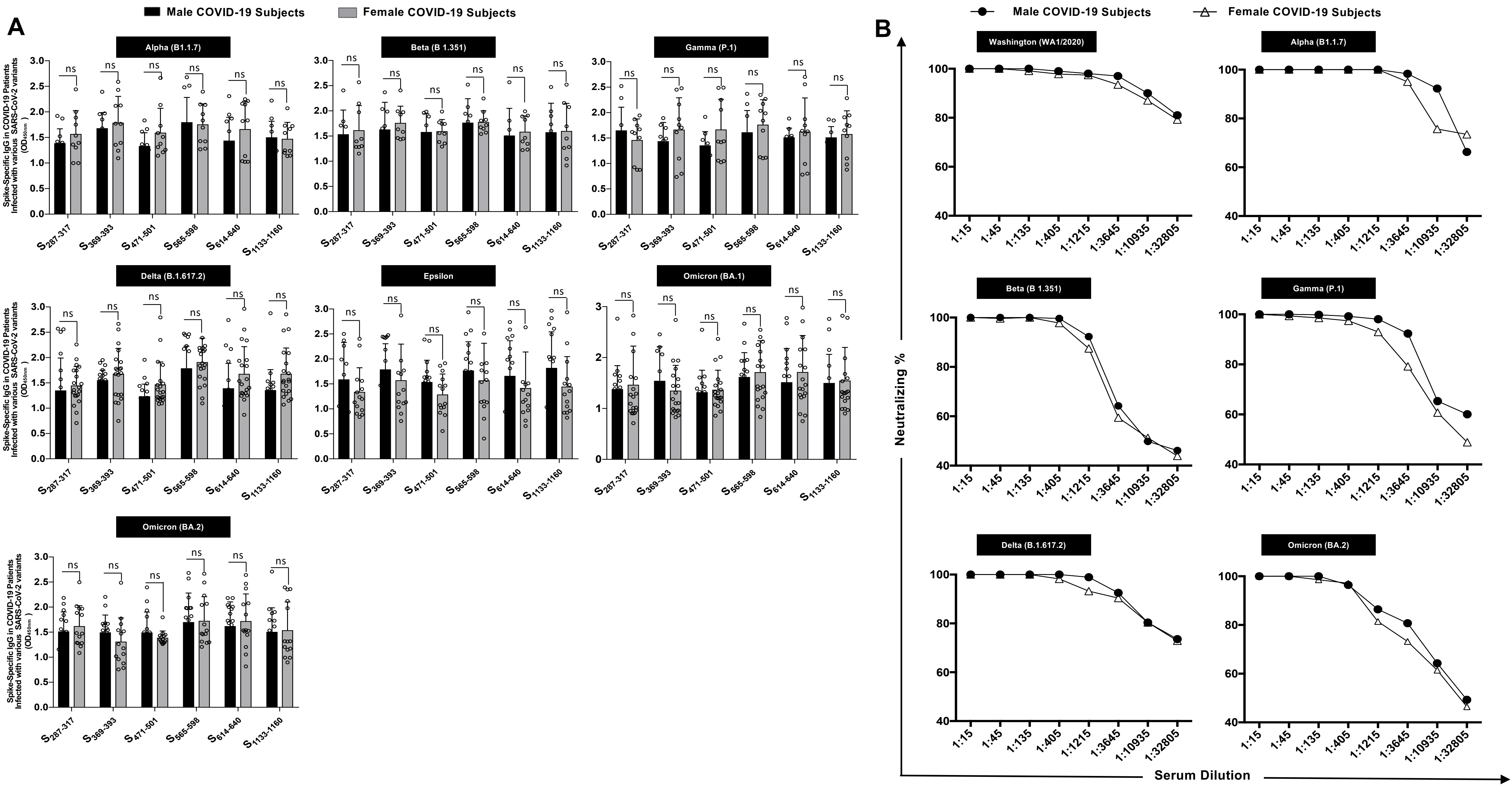

**Supplemental Figure S4. Gender-dependent immune responses against 'universal' B-cell epitopes in COVID-19 patients exposed to different SARS-CoV-2 variants of concern: Highly conserved COVID-19 peptide immunogenicity evaluation.** Similar to (**Fig. 2**) and (**Fig. 3** ), bar graphs illustrate the peptide binding IgG level for the 6 “universal” B cell epitopes measured by ELISA (*panel* **A**). Serum samples were collected from COVID-19 patients infected with various SARS-CoV-2 variants of concern (VOC) and categorized into two distinct groups based on gender. The (**B**) *panel* shows neutralization (%) by sera from these patients against the different SARS-CoV-2 VOCs Alpha (B.1.1.7), Beta (B.1.351), Epsilon (B.1.427/B.1.429), Delta (B.1.617.2), and Omicron (BA.2). Bars represent means ± SEM. Data were analyzed using student's t-test and multiple t-tests. Results were considered statistically significant at $P < 0.05$, with statistical correction applied using the Holm-Sidak method.

*Supplemental Table. 1*

| Spike protein position | Start position in sequence | End position in sequence | Sequence |
|---|---|---|---|
| $S_{287-317}$ | 287 | 317 | DAVDCALDPLSETKCTLKSFTVEKGIYQTSN |
| $S_{524-558}$ | 524 | 558 | VCGPKKSTNLVKNKCVNFNFNGLTGTGVLTESNKK |
| $S_{544-578}$ | 544 | 578 | NGLTGTGVLTESNKKFLPFQQFGRDIADTTDAVRD |
| $S_{565-598}$ | 565 | 598 | QFGRDIADTTDAVRDPQTLEILDITPCSFGGVSVI |
| $S_{601-628}$ | 601 | 628 | GTNTSNQVAVLYQDVNCTEVPVAIHADQ |
| $S_{614-640}$ | 614 | 640 | QDVNCTEVPVAIHADQLTPTWRVYSTGS |
| $S_{802-819}$ | 802 | 819 | FSQILPDPSKPSKRSFIE |
| $S_{888-909}$ | 888 | 909 | FGAGAALQIPFAMQMAYRFNGI |
| $S_{440-470}$ | 440 | 470 | NLDSKVGGNYNYLYRLFRKSNLKPFERDIST |
| $S_{456-485}$ | 456 | 485 | FRKSNLKPFERDISTEIYQAGSTPCNGVEG |
| $S_{471-501}$ | 471 | 501 | EIYQAGSTPCNGVEGFNCYFPLQSYGFQPTN |
| $S_{369-393}$ | 369 | 393 | YNSASFSTFKCYGVSPTKLNDLCFT |
| $S_{1133-1160}$ | 1133 | 1160 | VNNTVYDPLQPELDSFKEELDKYFKNHT |
| $S_{1145-1172}$ | 1145 | 1172 | LDSFKEELDKYFKNHTSPDVDLGDISGI |
| $S_{329-356}$ | 329 | 356 | FPNITNLCPFGEVFNATRFASVYAWNRK |
| $S_{338-363}$ | 338 | 363 | FGEVFNATRFASVYAWNRKRISNCVA |
| $S_{13-37}$ | 13 | 37 | SQCVNLTTRTQLPPAYTNSFTRGVY |

**Supplemental Table S1. The Amino acid sequences and selection criteria of B-cell epitopes from the spike protein of SARS-CoV-2.** The table presents the amino acid sequences of 17 B-cell epitopes derived from the spike protein of SARS-CoV-2. These epitopes were selected based on their high conservation across various coronaviruses, including SARS-CoV-2, the four major "common cold" coronaviruses (CoV-OC43, CoV-229E, CoV-HKU1, and CoV-NL63), and SARS-like coronaviruses (SL-CoVs) isolated from bats, civet cats, pangolins, and camels. Additionally, epitope selection considered the likelihood of each linear epitope being exposed on the surface of infected target cells.
